# Soft-metal(loid)s induce protein aggregation in *Escherichia coli*

**DOI:** 10.1101/2023.08.21.554180

**Authors:** Fabián A. Cornejo, Claudia M. Muñoz-Villagrán, Roberto A. Luraschi, María P. Sandoval-Díaz, Camila A. Cancino, Benoit Pugin, Eduardo H. Morales, Jeff S. Piotrowski, Juan M. Sandoval, Claudio C. Vásquez, Felipe A. Arenas

## Abstract

Metal(loid) salts have been used to treat infectious diseases due to their exceptional biocidal properties at low concentrations. However, the mechanism of their toxicity has yet to be fully elucidated. The production of reactive oxygen species (ROS) has been linked to the toxicity of soft metal(loid)s such as Ag(I), Au(III), As(III), Cd(II), Hg(II), and Te(IV). Nevertheless, few reports have described the direct, or ROS-independent, effects of some of these soft-metal(loid)s on bacteria, including the dismantling of iron-sulphur clusters [4Fe-4S] and the accumulation of porphyrin IX. Here, we used genome-wide genetic, proteomic, and biochemical approaches under anaerobic conditions to evaluate the direct mechanisms of toxicity of these metal(loid)s in *Escherichia coli*. We found that certain soft-metal(loid)s promote protein aggregation in a ROS-independent manner. This aggregation occurs during translation in the presence of Ag(I), Au(III), Hg(II), or Te(IV) and post-translationally in cells exposed to Cd(II) or As(III). We determined that aggregated proteins were involved in several essential biological processes that could lead to cell death. For instance, several enzymes involved in amino acid biosynthesis were aggregated after soft-metal(loid) exposure, disrupting intracellular amino acid concentration. We also propose a possible mechanism to explain how soft-metal(loid)s act as proteotoxic agents.

## 1. Introduction

Some metallic elements (e.g., sodium, iron, copper, and cobalt) are essential for life because their unique chemical properties are indispensable for cellular functions. These metals participate in redox reactions, provide structural stability, and enable critical cellular processes such as electron transfer and catalysis (Lemire et al., 2013). However, in excess, these metals can become toxic (Nies, 1999). On the other hand, non-essential metal(loid) ions such as mercury, arsenic, tellurium, and silver are highly toxic to most organisms even at micromolar concentrations (Nies, 1999; Taylor, 1999). Ions of these elements, due to their high polarizability, are classified as soft acids or soft-metal(loid)s. They tend to react with intracellular molecules containing soft bases like sulfhydryl, thioesters, phenyl, or imidazole groups, forming covalently bound adducts (Pearson, 1966; Lemire et al., 2013). Conversely, metals such as Mg(II), Na(I), and K(I) are classified as hard metals and primarily interact through ionic bonds with hard bases like sulfate, carboxylate, phosphate, and amine groups (Pearson, 1966; Lemire et al., 2013).

Soft-metal(loid)s toxify cells by producing Reactive Oxygen Species (ROS) and inducing oxidative stress damage (Parvatiyar et al., 2005; Pérez et al., 2007; Tremaroli et al., 2007; Pacheco et al., 2008; Park et al., 2009; Lemire et al., 2013; Muñoz-Villagrán et al., 2020). Some metal(loid)s can produce ROS directly via Fenton chemistry [e.g., Fe(II) and Cu(I)], or indirectly, where Fenton-inactive metal(oid)s such as As(III), Cd(II), and Te(IV) deplete reduced glutathione, compromising the redox state and oxidative stress response of the cell (Liochev. and Fridovich, 1999; Turner et al., 2001; Valko et al., 2005). The depletion of major cellular sulfhydryl reserves seems to be a critical indirect mechanism for oxidative stress induced by metal(loid)s (Bagchi and Stohs, 1993). The consequences of cells under oxidative stress include various dysfunctions mediated by direct damage to lipids, proteins, and DNA (Ercal et al., 2001). Metals like Cd(II), Cu(I), Hg(II), Te(IV), and Ni(II) can trigger lipid peroxidation in different organisms (Company et al., 2004; Paraszkiewicz et al., 2010; Pradenas et al., 2013). Membrane damage can be caused by Cu(II) or Cd(II), and Ag(I) can cause a loss of membrane potential (Dibrov et al., 2002; Hong et al., 2012). Double-stranded DNA can be damaged by oxidation, thereby inducing cell death and mutagenicity (Ercal et al., 2001), as demonstrated with iron (Nunoshiba et al., 1999) and via genotoxicity assays with Mn, Cr, Cd, and other metals (Lemire et al., 2013). ROS can also induce protein oxidation and dysfunction, forming carbonyl derivatives through the metal-catalyzed oxidation of several amino acid side chains (such as histidine, arginine, lysine, and proline). Hence, carbonyl group levels are often used as a marker of oxidative protein damage (Stadtman and Levine, 2003). In other instances, metal(loid)-induced ROS dismantle Fe-S clusters in some dehydratases (Imlay, 2006; Calderón et al., 2009; Gomez et al., 2014).

The soft or hard acid nature of a specific metal(loid) can explain its toxicity and reactivity toward soft-base ligands within the protein matrix (Medici et al., 2021; Peana et al., 2021). For instance, Ni(II) can replace the structural Zn(II) present in the metal-binding sites of some proteins. These metal-binding sites can be exchanged by other more competitive divalent soft-metals, resulting in mismetallation and protein function loss (Macomber et al., 2011; Quintal et al., 2011). Certain metal(loid)s can damage enzymes that display reactive Cys residues at their active sites (Rajanna et al., 1990; Fadeeva et al., 2011). The soft-metal(loid)s Cu(I), Hg(II), Ag(I), Cd(II), and Te(IV) can dismantle solvent-exposed [4Fe-4S] clusters of *Escherichia coli* aconitases (Macomber and Imlay 2009; Calderón et al., 2009; Xu and Imlay 2012), releasing Fenton-active iron into the cytoplasm, which in turn generates ROS.

Despite the similar chemical reactivity of soft-metal(loid)s, few studies have identified cell targets beyond ROS-mediated oxidative stress. In this study, we conducted a genome-wide screening of an *E. coli* deletion collection challenged with soft-metal(loid)s under anaerobic conditions to identify ROS-independent targets. Strains lacking genes involved in protein homeostasis showed reduced fitness under these conditions, suggesting that proteins aggregated when exposed to Ag(I), Au(III), As(III), Cd(II), Hg(II), or Te(IV). Protein aggregation induced by Ag(I), Au(III), Hg(II), and Te(IV) required active translation, whereas As(III) and Cd(II) do not, suggesting that the mechanism by which these metals act during protein aggregation is different.. In summary, we demonstrated that, in a ROS-independent manner, soft-metal(loid)s act as proteotoxic agents, leading to the accumulation of aggregated proteins, most likely due to at least two distinct mechanisms.

## 2. Methods

### 2.1 Bacterial strains

*E. coli* K-12 BW25113 (National Institute of Genetics, Microbial Genetics Laboratory, NBRP, Japan) was used as the model strain. Unless otherwise indicated, all cultures were grown at 37 °C in an anaerobic chamber (Coy Laboratory Products Inc., 100% N_2_ atmosphere) with constant shaking (150 rpm) in MOPS minimal medium supplemented with 0.2 % (w/v) glucose. The barcoded deletion collection was derived from *E. coli* K-12 BW38028 (Otsuka et al., 2015).

### 2.2 Chemical genomic profiling of soft-metal(loid)s in *E. coli*

The pooled *E. coli* deletion collection was independently inoculated in media supplemented with the following soft-metal(loid) salts (resuspended in H_2_O) at concentrations that resulted in a 20-30% reduction of OD_600_ compared to unexposed controls after 24 h: AgNO_3_ (625 nM), HAuCl_4_ (2 µM), NaAsO_2_ (125 µM), CdCl_2_ (31.25 µM), HgCl_2_ (375 nM) or K_2_TeO_3_ (14.7 µM). For control experiments, salts were replaced with H_2_O. Each pooled competition experiment was conducted in 200 µL cultures in triplicate at 37 °C for 24 h. Genomic DNA was extracted using the Wizard Genomic DNA Purification Kit (Promega, Cat. No. A1120). Strain-specific barcodes from each culture were amplified using indexed primers designed for multiplexed Illumina sequencing. The forward primer contained the Illumina-specific P5 sequence, a 10 bp index tag (x’s), and the 19 bp *E. coli* deletion collection common priming site: 5’
s-AATGATACGGCGACCACCGAGATCTACACTCTTTCCCTACACGACGCTCTTCCGATCT xxxxxxxxxxAATCTTCGGTAGTCCAGCG-3′. The reverse primer included the Illumina-specific P7 sequence and 20 bp *E. coli* common priming site: 5′-CAAGCAGAAGACGGCATACGAGCTCTTCCGATCTTGTAGGCTGGAGCTGCTTCG-3′. As previously described, barcodes were amplified by PCR, pooled, gel-purified, and quantified by quantitative PCR (Piotrowski et al., 2015). For barcode sequencing, samples were run on an Illumina HiSeq2500 in rapid run mode for 50 cycles at a loading concentration of 15 pM. The resulting fastq file was used for the analysis of sensitive and resistant mutants. Normalized counts were compared to a control solvent (water) to identify compound-specific responses among gene deletion mutants (chemical-genetic interaction score).

Sequence data were processed using BEANcounter (Simpkins et al., 2019) and EdgeR (Robinson et al., 2010). The chemical-genetic interaction score was computed as z-scores, representing the standardized deviation of each strain in treatments compared to their counterpart strain in the control solvent, for the 3551 mutants quantified (Piotrowski et al., 2017; Simpkins et al., 2019). CG-scores were analyzed using the DAVID database (Huang et al., 2009). Gene functional annotations were retrieved from EcoCyc (Keseler et al. 2017).

### 2.3 Isolation of aggregated proteins

Protein aggregation was assessed using the method described by Tomoyasu et al. (2001). Briefly, saturated cultures (16 h) were inoculated into 250 ml flasks containing 60 mL of MOPS medium, to a starting OD_600_ of approximately 0.05, and grown anaerobically at 37 °C with constant shaking until they reached an OD_600_ of approximately 0.3. Cultures were individually exposed to soft-metal(loid) concentrations (as indicated in each figure) and incubated for 2 h at 37 °C. Fifty mL of each treated culture were centrifuged at 5,000 x g for 10 min at 4 °C. The resulting cell pellets were suspended in 250 µL of 10 mM potassium phosphate buffer pH 6.5 containing 1 mM EDTA (buffer A), supplemented with 20 % (w/v) sucrose and 1 mg/mL lysozyme, and incubated on ice for 30 min. Then, 360 µL of buffer A was added, and cells were disrupted by sonication on ice (eight 15-s cycles with 45-s rests at 60 % amplitude). Cell debris was removed by centrifugation at 2,000 x g for 15 min at 4 °C. Both aggregated and membrane proteins were centrifuged at 15,000 x g for 20 min at 4 °C and frozen at -80 °C for later processing. Sedimented proteins were suspended in 400 µL of buffer A by brief sonication (one pulse, 5 s at 60% amplitude) and sedimented again at 15,000 x g for 20 min at 4 °C. Membrane proteins were removed by suspending the pellet via brief sonication (one pulse, 5 s at 60% amplitude) in 320 µL of buffer A containing 2 % (v/v) NP-40 (Abcam, Cat. No. ab142227). The aggregated proteins were sedimented at 15,000 x g for 30 min at 4 °C. This washing procedure was repeated four times. The NP-40 insoluble pellet was rinsed with 400 µL of buffer A and finally suspended in 200 µL of the same buffer. Aggregated proteins were quantified by the Bradford method (Bradford, 1976), resolved by SDS-PAGE (12 %), and visualized by silver staining.

### 2.4 Translation-arrested cells and pulse-chase of aggregated proteins

For experiments involving translation arrest, 100 µg/mL of chloramphenicol (CHL) was added to the cultures five min prior to the soft-metal(loid) treatment. For pulse-chase experiments, cultures (with an OD_600_ of approximately 0.3) were pulse-labeled with 250 µM 4-azido-L-homoalanine (AHA, Jena Bioscience, Cat. No. CLK-AA005) and incubated for 15 min at 37 °C. The chase was initiated by adding L-methionine (Sigma-Aldrich, Cat. No. M9625) to a final concentration of 250 µM and incubated for 30 min at 37 °C. Cells were then exposed to soft-metal(loid)s at the indicated concentrations for 2 h at the same temperature. Aggregated proteins were isolated as described above, but EDTA was omitted from the buffers to prevent Cu chelation in subsequent reactions. Protein aggregates were washed with phosphate saline buffer (PBS) and suspended through a brief (5-s) sonication in 200 µL of Click reaction buffer. This buffer contained 50 µM acetylene-PEG4-Biotin (Jena Bioscience, Cat. No. CLK-TA105), 1 mM TCEP (Tris(2-carboxyethyl)phosphine hydrochloride Sigma-Aldrich, Cat. No. C4706), 100 µM THPTA (Tris(3-hydroxypropyltriazolylmethyl)amine, Jena Bioscience, Cat. No. CLK-1010), and 1 mM CuSO_4_ (Sigma-Aldrich, Cat. No. C8027) in PBS, and incubated for 30 min at 37 °C. Biotinylated protein aggregates were washed twice with PBS and then analyzed via Western blotting.

### 2.5 Western blotting of biotinylated aggregated proteins

Five hundred nanograms of aggregated proteins were fractionated using SDS-PAGE (12%). Proteins were then transferred to PVDF membranes (Bio-Rad, Cat. No. 1620177) at 40 mA overnight at 4 °C. The membranes were blocked for 1 h in a Tris-buffered saline (TBS) buffer that was supplemented with 1 % (v/v) Tween-20 (TBS-T) and 3 % (w/v) BSA. After an overnight incubation at 4 °C with a 1:10,000 dilution of Streptavidin-HRP (Sigma-Aldrich, Cat. No. S5512) in TBS-T plus 3% (w/v) BSA, the biotinylated proteins were visualized using SuperSignal Western blot FEMTO substrate (Thermo Scientific, Cat. No. 34094).

### 2.6 Proteomic profiling of aggregated proteins

Samples of aggregated proteins, obtained after a 2 h exposure to metal(loid) concentrations affecting the cell viability in a similar extend (10 µM AgNO_3_, 5 µM HAuCl_4_, 200 mM NaAsO_2_, 500 µM CdCl_2_, 5 µM HgCl_2_, or 20 µM K_2_TeO_3)_, were flash-frozen in an ethanol dry-ice bath and processed for label-free quantification at Bioproximity in Virginia, USA. Only proteins identified with at least two unique peptides were included in the analysis. All proteins identified in the control sample were excluded from the analysis, mainly ribosomal proteins and translation factors. Functional protein associations and enrichment analysis were evaluated using the stringApp in Cytoscape (Doncheva et al., 2018). Enrichment analysis of aggregated proteins in each treatment was conducted using the DAVID database (Huang et al., 2009). All properties and subcellular locations of *E. coli* proteins were retrieved from EcoCyc (Keseler et al., 2017).

### 2.7 Cell viability

Cell viability was determined by diluting 20 µL of each treatment with 180 µL of a sterilized 0.9 % (w/v) NaCl solution. After diluting up to 10^−7^, 4 µL of each dilution was spotted on LB (Luria Bertani broth) agar plates and incubated overnight at 37 °C in anaerobic conditions.

### 2.8 Amino acids extraction and quantification

Amino acids were extracted using the method described by Steinfeld et al. (2014). Briefly, 60 mL of anaerobic *E. coli* cultures at OD_600_ 0.3 in MOPS media were treated for 2 h with defined concentrations of soft-metal(loid)s. Twenty mL from each treatment were centrifuged at 4,000 x g for 10 min at 4 °C. Cell pellets were resuspended in 1 mL of ice-cold MOPS media and centrifuged as described above. Then, cells were lysed with 1 mL of 1 N HCl, incubated for 5 min at room temperature, and centrifuged at 19,000 x g for 20 min at 20 °C. The supernatant was dried in a Vacuum Concentrator SpeedVac SPD120 for 3 h at 50 °C and 1.5 h at 35 °C. The resulting pellets were flash-frozen at -80 °C until use. Pellets were resuspended in 300 µL of water by sonication for 5 s, and two consecutive extractions with 500 and 300 µL of chloroform were performed. The aqueous layers were combined, filtered through a 0.22 µm PVDF filter (4 mm), and dried for 80 min at 65 °C and 80 min at 55 °C. The pellets were kept flash-frozen at -80 °C until used.

Amino acids were quantified using diethyl ethoxymethylenemalonate (DEEMM) derivatization followed by UPLC-DAD quantification as described in Otaru et al., 2021, with few modifications. Each pellet was resuspended in 50 µL of 0.1 N HCl. After dissolution, 87.5 µL of 1 M borate buffer pH 9.0, 37.5 μL methanol, 2 μL of 2 g/L L-2-aminoadipic acid (in 0.1 M HCl; internal standard), and 1.75 μL DEEMM were added. Analytes were derivatized for 45 min at room temperature in an ultrasonic water bath, and the samples were then heated for 2 h at 70 °C to stop the reaction. After that, samples were filtered (0.45 µm membrane) and transferred into glass vials for analysis.

Quantification was carried out using an H-Class Acquity UPLC system (Waters Corp., Milford, MA, USA) equipped with a photodiode array detector (DAD). The derivatized molecules were separated using a gradient of (A) 25 mM acetate buffer pH 6.6, (B) methanol, and (C) acetonitrile in a Waters Acquity UPLC BEH C18 1.7 µm column (2.1 x 100 mm) at 40 °C, as described in Otaru et al., (2021). One µL sample was applied to the column and eluted at a flow rate of 0.46 mL/min. Individual compounds were quantified at 280 nm using the internal standard method. Data processing was performed using Empower 3 software (Waters).

### 2.9 Statistical analysis

Biochemical assays were repeated at least three times, and the data were represented as bar plots (mean with standard deviation or standard error), scatter plots (mean with standard error), or heatmaps (mean). In the case of SDS-PAGE and Western blot, representative gels were shown.

The DAVID platform was used to compute enrichment analysis statistics. Two-sample student’s *t*-tests, ANOVA, post hoc tests, and normal distribution statistics were calculated using R.

## 3. Results and Discussion

### 3.1 Chemical genomic analysis of soft-metal(loid) toxicity in *E. coli*

The toxicity of metal(loid)s in bacteria has been extensively studied and has been mostly related to the production of ROS. These highly reactive molecules rapidly oxidize several macromolecules, including DNA, membranes, and proteins. Moreover, protein aggregation was observed in yeast exposed to As(III) or Cd(II) under aerobic conditions (Jacobson et al., 2012; Jacobson et al., 2017). As(III) and Cd(II) displayed mutagenic activities dependent on the presence of RecA, suggesting single or double breaks in the DNA molecule (Lemire et al., 2013). However, the presence of oxygen in these experiments makes it difficult to determine whether the observed effect is a direct result of metal(loid) damage or ROS production. To directly identify the direct intracellular targets of soft-metal(loid)s, we challenged *E. coli* cells under anaerobic conditions, where ROS cannot be formed.

We conducted chemical-genomic profiling of *E. coli* to identify deletions responsive to the soft-metal(loid)s Ag(I), Au(III), As(III), Cd(II), Hg(II), and Te(IV) under anaerobic conditions by using a published barcoded deletion collection (Otsuka et al., 2015; Morales et al., 2017). Each mutant in this barcoded collection has a unique 20-nucleotide barcode that allows quantification of its abundance through next-generation sequencing in a pooled competition experiment. We challenged the pooled collection with soft-metal(loid) concentrations that produced a mild (20–30%) reduction of the optical density (OD_600_) after 24 h compared to the non-treated control to avoid depletion of mutants involved in the general stress response. The chemical-genetic score (from now on, CG-scores, see **Table S1**) of each knockout strain served as an indicator of the relative abundance after the challenge (Simpkins et al., 2019).

Deletion of genes involved in fundamental processes such as cell division, lipopolysaccharide core region synthesis, protein import, homologous recombination, conjugation, and cell shape regulation, was detrimental to soft-metal(loid) exposure (**Figure S1A**). These results suggest that soft-metal(loid)s have a pleiotropic effect and affect multiple molecular targets, which ultimately leads to cell death. Strikingly, our analyses revealed that mutants in genes encoding proteins required for homologous recombination-DNA repair (Figure 1A and S1A) and protein aggregation (**Figure 1A**), were among the most impacted by toxic exposure, strongly suggesting that soft-metal(loid)s could damage DNA and proteins under anoxic conditions.

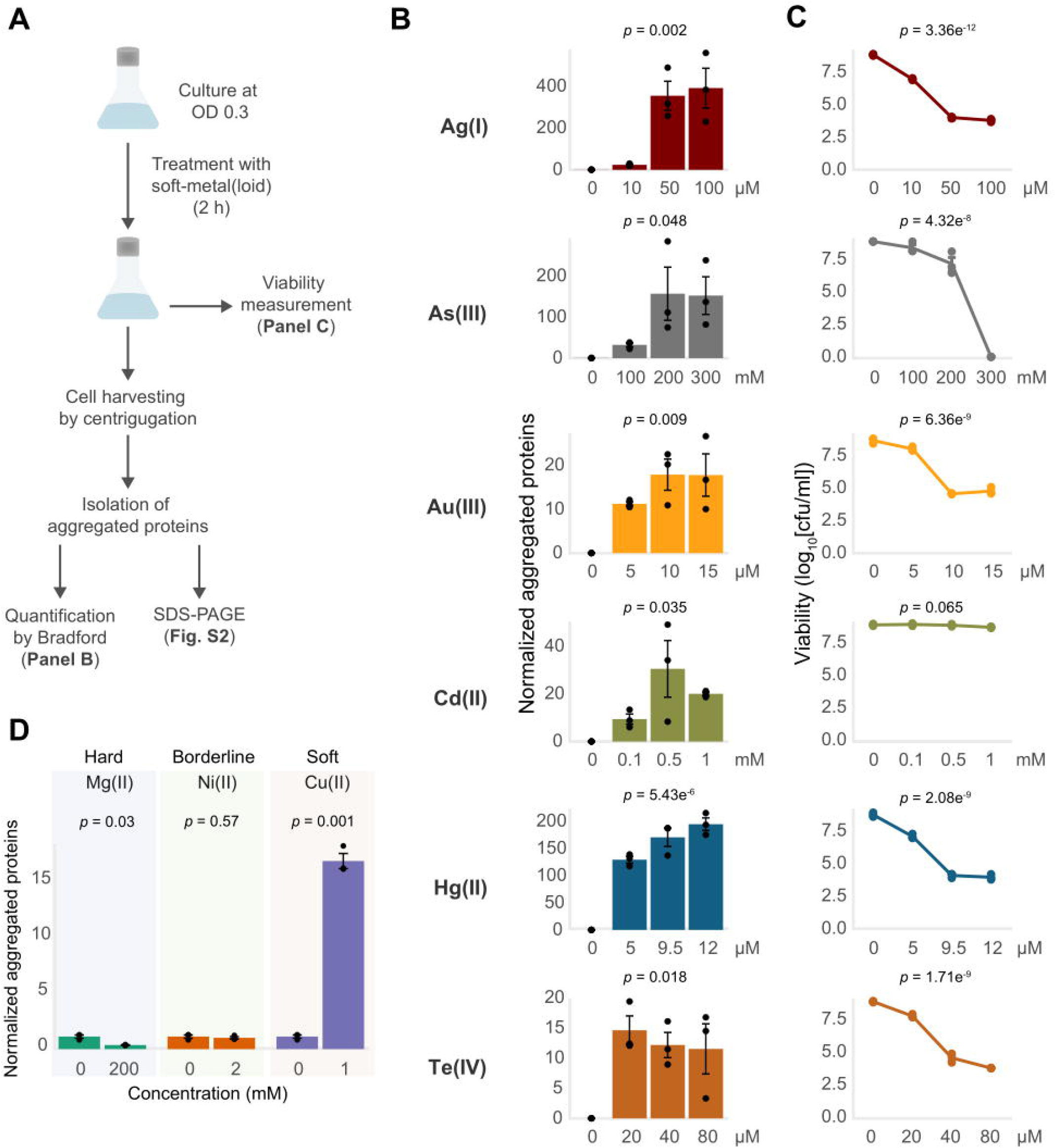

### 3.2 DNA damage produced by Au(III)

Of all the tested soft-metal(loid)s, Au(III) had the greatest effect on the fitness of mutants in genes involved the homologous recombination DNA repair pathways such as *recABC* and *ruvABC*, among others (Figure S1B), suggesting that Au(III) could induce DNA breaks independently of ROS. Based on this, we decided to use Au(III) as a model to investigate this effect. Supporting our hypothesis, we detected free 3’OH ends by TUNEL-assay after exposure to 10 µM HAuCl_4_ for 30 min (**Figure S1C**). Although this assay detects both single-strand (SSB) and double-strand breaks (DSB), the deletion of genes involved in the RecBCD pathway suggests that the later might be occurring.

Au(III), and perhaps also other soft-metalloids, seems to generate DSBs by a different mechanism than other molecules already studied. For instance, quinolones inhibit the topoisomerases II and IV in the ligation process, generating DSB (Kohanski, Dwyer and Collins, 2010). Cd(II) has been described as a potential topoisomerase II inhibitor by binding to critical Cys residues. However, this inhibition completely disrupts its DNA-cleaving activity (Wu, Yalowich and Hasinoff, 2011). Other topoisomerase II inhibitors that react with free Cys do not generate DSBs (Jensen et al., 2002; Wu et al., 2008; Sciandrello et al., 2010). This suggests that a plausible reaction of Au(III) with the enzyme’s Cys residues cannot lead to the DSB observed in ROS-independent conditions.

### 3.3 Soft-metal(loid)s induce protein aggregation

Previous research has suggested that soft metal(loid)s can induce protein aggregation and damage proteins under aerobic conditions. A prominent example of this is the interaction of Hg(II) and Cd(II) with luciferase during an *in vitro* refolding process, leading to its inactivation (Sharma et al., 2008). A recent study discovered that the soft-metal Cu(I) can cause protein aggregation in *E. coli* independently of ROS (Zuily et al., 2022).

Our analysis showed that deletion mutants in genes implicated in the protein quality control (PQC) network had decreased fitness in *E. coli* exposed to soft-metal(loid)s, suggesting these metal(loid)s can induce protein misfolding and aggregation in a ROS-independent manner (**Figure 1A**).

Under standard conditions, misfolded proteins can be refolded by chaperones or degraded by various proteolytic systems, preventing their accumulation. However, the protein quality control network can be overwhelmed under stress conditions, leading to an accumulation of protein aggregates (Tyedmers et al., 2010). Once these aggregated proteins are formed in *E. coli*, they can be refolded by ClpB (assisted by DnaK, DnaJ, and GrpE) or degraded by AAA+ proteases like ClpAP, HslUV, or Lon (Dougan et al., 2002; Haslberger et al., 2007; Tyedmers et al., 2010). DegS and DegP are two critical proteases involved in detecting misfolded and aggregated proteins in the periplasmic compartment. DegS functions as a regulatory protease that detects misfolded proteins through its PDZ domain and catalyzes the proteolysis of the anti-sigma factor RseA (Wilken et al. 2004) (**Figure 1B, right panel**). This process releases the alternative sigma factor RpoE, initiating a transcriptional response to manage misfolded proteins in the cytoplasmic and periplasmic compartments (Erickson and Gross 1989; Danese and Silhavy 1997). Egler et al. (2005) found that a Δ*rpoE* strain is sensitive to Cu(II), Cd(II), and Zn(II), but the authors did not speculate about a potential metal-mediated proteotoxic effect. DegP is a RpoE-induced serine protease/chaperone that degrades misfolded proteins in the periplasm. In our analysis, mutants lacking cytoplasmic and periplasmic proteases and chaperones, such as DegP, DegQ, the AAA+ protease ClpAP, and its adaptor ClpS (Dougan et al., 2002), displayed negative fitness when exposed to soft-metal(loid)s (**Figure 1B**). A similar negative fitness was observed for mutants lacking the refoldase ClpB and other cytoplasmic chaperones like DnaK and the small heat shock proteins IbpA and IbpB, all of which form an intricate network for protein disaggregation (Tyedmers et al., 2010). Consistent with this, previous transcriptomic analyses of *E. coli* exposed to Cd(II), Hg(II), and Te(IV) showed the induction of chaperone and protease systems such as ClpB, DnaK, and ClpP, among others (Wang and Crowley 2005; Molina-Quiroz et al. 2014; LaVoie and Summers 2018), supporting a proteotoxic effect of these metal(loid)s.

An intriguing observation was that Lon deletion mutants showed improved fitness following soft-metal(loid) treatment. Lon is one of the main AAA+ proteases involved in the degradation of misfolded proteins, thereby preventing aggregation (Rosen et al., 2002). Some researchers have reported that it plays a minor role in protein aggregate clearance (Vera et al., 2005), but this doesn’t explain the increased resistance to soft-metal(loid)s. This phenotype could result from Lon-dependent regulatory proteolysis of an unknown factor that when stabilized, could help combat soft-metal(loid) toxicity. Further studies are required to elucidate the role of Lon in this process.

To directly test if soft-metal(loid)s caused protein misfolding and aggregation under anaerobic conditions, we challenged exponentially growing wild type *E. coli* BW25113 to increasing concentrations of soft-metal(loid)s for 2 h and immediately quantified aggregated proteins (**Figure 2A-C, Figure S2**). For all soft-metal(loid)s tested, treatment led to protein aggregation in a concentration-dependent manner and started even at low toxic concentrations (**Figure 2B and S2A)**.

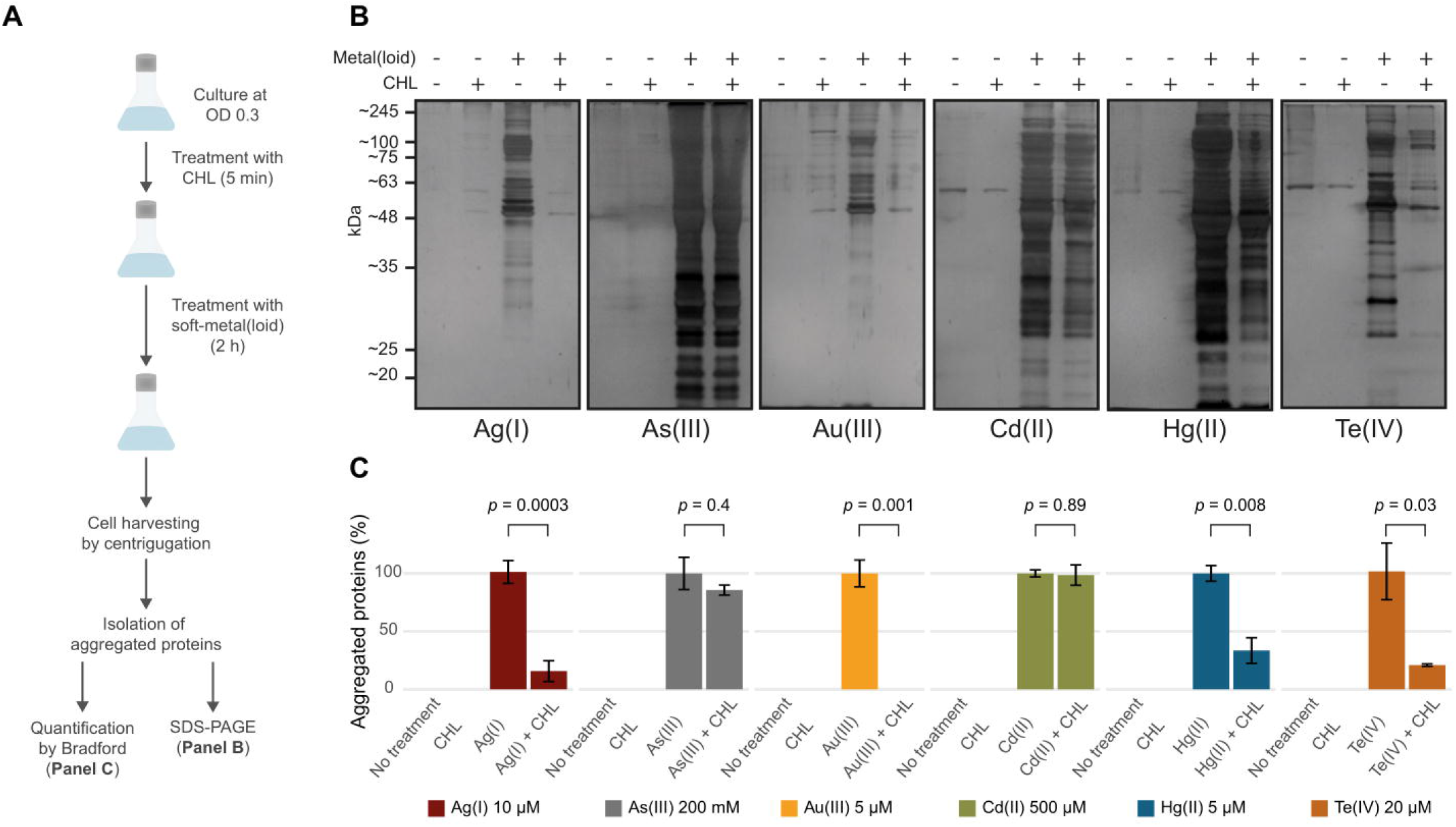

Interestingly, while all the tested soft-metal(loid)s caused protein aggregation, their effects on cell viability varied considerably (**Figure 2B-C**). For instance, 9.5 µM Hg(II) induced approximately 150 µg of aggregated proteins per mg of total protein, significantly decreasing viability by 5 orders of magnitude. In contrast, 200 mM As(III) produced similar number of aggregated proteins to Hg(II), but its impact on cell viability was noticeably less (**Figure 2B-C**). Others, like Te(IV) and Au(III), strongly affected viability at 20 and 10 µM, respectively; however, they induced less protein aggregation compared to Ag(I), Hg(II), and As(III). This indicates that for each metal(loid) there is no direct correlation between its effect on viability and the amount of protein aggregates (**Figure 2B-C**).

Recently, Zuily et al. (2022) demonstrated that the soft-metal Cu(I), and to a lesser extent Cu(II), can cause protein aggregation independent of ROS. In agreement with these results, we observed protein aggregation induced by Cu(II) at millimolar concentrations (0.5 to 2.0 mM), comparable to protein aggregation induced by some other soft-metal(loid)s tested (**Figure 2D and S2B**). Altogether, this suggests that the soft-acid property of these metal(loid)s could explain their reactivity towards proteins and cause aggregation. To test this, we exposed *E. coli* to the hard-acid metal Mg(II) and the borderline metal Ni(II) (**Figure 2D and S2B**). Exposure to Mg(II) or Ni(II) in the millimolar range did not result in protein aggregation, suggesting that the soft-acid property, and thus the covalent nature of the metal-ligand interaction, might be important for inducing protein aggregation.

We next sought to assess whether protein aggregation could happen during or required active translation, as nascent proteins are not fully folded. The exposed soft residues may interact with these metal(loid)s through covalent bond formation, leading to unexpected geometries that could cause protein misfolding. To test this, we halted active translation by using chloramphenicol (CHL) for 5 min prior to toxic exposure and measured aggregated proteins (**Figure 3**). CHL pre-treatment did not affect protein aggregation mediated by As(III) and Cd(II), suggesting that it does not depend on active translation (**Figure 3B-C**). Conversely, Au(III)-induced aggregation was completely inhibited when the translation was blocked (**Figure 3B-C**), while for. Ag(I), Hg(II), and Te(IV) it was inhibited by 85%, 65%, and 80%, respectively (**Figure 3C**). We hypothesize that the metal(loid) Ag(I), Hg(II), Au(III), and Te(IV) primarily cause protein aggregation by interacting with nascent proteins, while Cd(II) or As(III) can also interact with pre-folded proteins and produce aggregation by y different mechanism. To directly test this, we conducted pulse-chase experiments in cells treated with the metalloids As(III) and Te(IV) as models. Proteins were pulse-labelled with 4-azido-L-homoalanine (AHA) for 15 min and chased with excess L-Met for 30 min to produce labeled folded proteins and unlabeled nascent proteins during toxicant exposure (**Figure S3**). As hypothesized, the label was only detected in aggregates from cells treated with As(III), confirming that this oxyanion causes aggregation of proteins that are already folded, while Te(IV) appears to target nascent proteins (**Figure S3**).

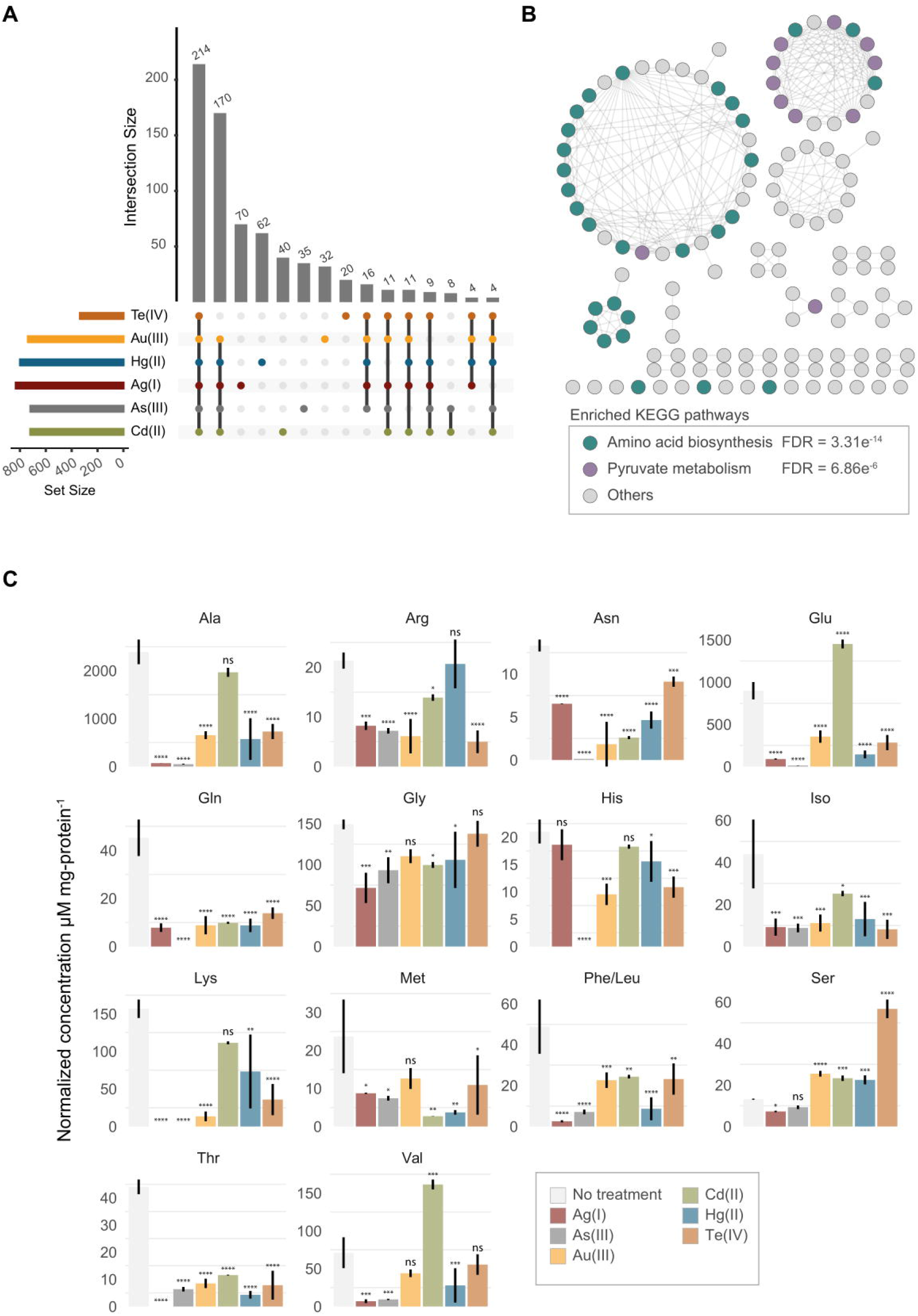

In summary, our results show that exposure to soft-metal(loid)s caused protein aggregation in a ROS-independent manner. In *E. coli* soft-metal(loid)s could cause protein aggregation through at least two hypothetical distinct mechanisms: (i) a translational-dependent mechanism where Hg(II), Au(III), Ag(I), and Te(IV) react with nascent proteins causing misfolding, and (ii) a translational-independent (or post-translational) one, where As(III) and Cd(II) preferentially target mature proteins. Alternatively, the latter could be attributed to the inhibition of disaggregation or protein degradation pathways by Cd(II) and As(III), which might ultimately lead to an imbalance in proteostasis and an accumulation of protein aggregates. In this context, Cd(II), Hg(II), and Pb(II) have been found to inhibit DnaK-, DnaJ-, and GrpE-assisted refolding and disaggregation in vitro (Sharma, Goloubinoff and Christen, 2008).

Protein aggregation at the translational level can be explained by the interaction between Hg(II), Au(III), Ag(I), and Te(IV) with soft-side chains such as those from Cys, His, Met, and Phe residues (Stadtman and Levine, 2003). This may hinder proper folding by introducing different coordination geometries and impacting folding rates, leading to misfolding, aggregation, and the sequestration of other proteins. Proteins likely react with these metal(loid)s through metal binding sites as they are being synthesized, resulting in incorrect metallation and misfolding. For instance, the Cu, Zn superoxide dismutase (Cu, Zn - SOD) from *S. cerevisiae* undergoes the exchange of Cu(I) for Ag(I) when the yeast is exposed to silver nitrate, generating an inactive Ag, Zn - SOD. Interestingly, Ag, Zn - SOD is less immunoreactive to conformational antibodies against Cu, Zn - SOD, suggesting a change in the protein’s folding or conformation (Ciriolo et al., 1994).

### 3.4 Identification of aggregated proteins

The reduction of soluble proteins and their diversion to the insoluble fraction may account for the toxicity of protein aggregates, consequently impacting normal metabolism and appropriate cell development (Mogk et al., 2018). To gain further insights on how this process impacts the resistance of *E. coli* against soft-metal(loid)s, we identified the aggregated proteins derived from the exposure to each toxic by mass spectrometry (Label-Free Quantification) (**Figure 4, Table S2**).

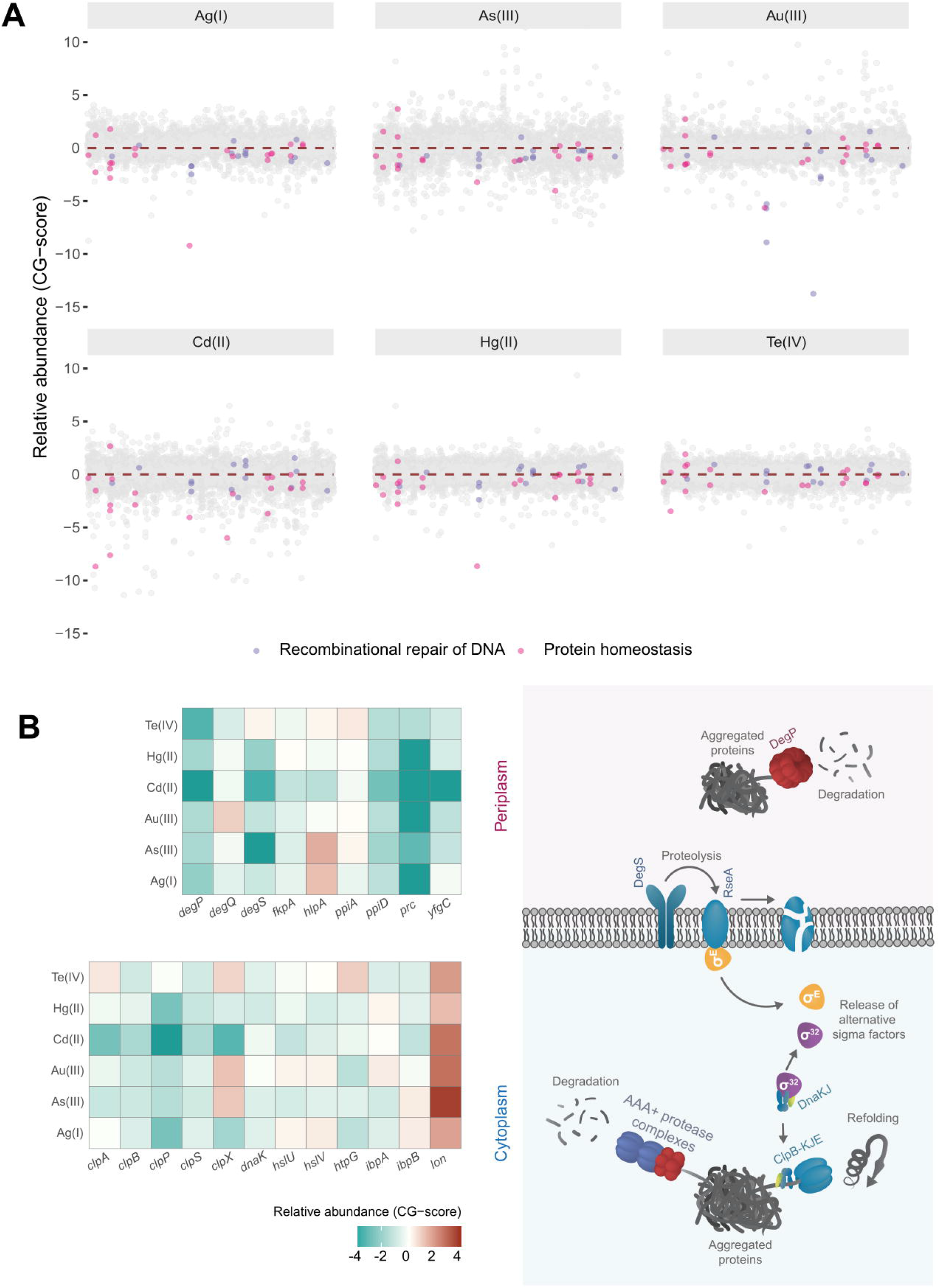

After treatment with Te(IV), Au(III), As(III), Hg(II), Cd(II), or Ag(I), we identified 335, 737, 718, 798, 719, and 831 insoluble proteins, respectively (**Figure 4A**). This represents between 7.5 and 18.6% of the proteins encoded by *E. coli*. Although the removal of periplasmatic chaperones and proteases resulted in decreased fitness after toxic exposure, the aggregated proteins that we identified are mainly cytoplasmic. For example, 78.5, 85.7, 87.2, 82.8, 88.9, and 87.1% of protein aggregates induced by Te(IV), Au(III), As(III), Ag(I), Cd(II), and Hg(II), respectively, were cytoplasmic proteins.

Next, to assess the presence of essential proteins within the aggregated material, we compared sets of aggregated proteins in each treatment to essential genes identified for a genome-wide deletion collection in MOPS media (Baba et al., 2006). The number of essential proteins in aggregates varied depending on the metal(loid) treatment: aggregates induced by Ag(I), Te(IV), As(III), Au(III), Hg(II), and Cd(II) contained 110, 21, 98, 102, 109, and 96 essential proteins, respectively. This indicates that protein aggregation induced by soft metal(loid)s disrupts crucial cellular processes by decreasing the soluble levels of essential proteins, leading to cell death. An interesting documented example of this phenomenon is MetA from *E. coli*, an essential thermosensitive protein in minimal media; depletion of this protein from soluble media due to high temperature-dependent aggregation causes a decrease in L-Met biosynthesis, impacting cell growth (Gur et al., 2002).

Among the total number of proteins identified in the aggregates induced by soft metal(loid)s, 214 were shared by all treatments (Figure 4A). A KEGG pathway enrichment analysis revealed that these proteins are primarily involved in metabolic pathways such as amino acid biosynthesis, pyruvate metabolism, central metabolism, and the biosynthesis of secondary metabolites (**Figure 4B, Table S3**). When proteins identified in every soft-metal(loid) treatment are examined individually, they participate in the same cellular processes and metabolic pathways as the aforementioned 214 proteins (**Figure 4A-B**). Surprisingly, Cd(II) and As(III) influenced the same processes as the other soft-metal(loid)s, despite inducing aggregation through a potentially different mechanism. Nonetheless, in our analysis, it is challenging to determine which proteins were directly misfolded by the metal(loid) treatment and which were subsequently trapped by the aggregates.

### 3.5 Soft-metal(loid)s perturb intracellular amino acid concentration

Since proteins involved in amino acid biosynthesis were among the most represented in protein aggregates for all treatments (**Figure 4B**), we speculated that amino acid levels might be affected after toxicant exposure. We measured the intracellular concentration of 15 amino acids in exponentially growing *E. coli* exposed to soft-metal(loid)s under the same conditions as in the protein identification analysis. Peaks for phenylalanine and leucine were not resolved, and their concentration represents the sum of both amino acids. Although none of the metal(loid)s depleted all amino acids, they impacted the levels of a significant number of them, ranging from 8 [Cd(II)] to 14 [Ag(I) and As(III)]. All soft-metal(loid) treatments decreased the intracellular concentration of asparagine, glutamine, isoleucine, phenylalanine + leucine, and threonine (**Figure 4C**). For asparagine, isoleucine, and leucine, we detected proteins involved in their biosynthesis in the aggregates of most of the soft-metal treatments (**Figure 4C** and **Table S2**). AsnA (aspartate-ammonia ligase (ADP-forming) (EC 6.3.1.4), and IlvA (threonine ammonia-lyase, EC 4.3.1.19) have been previously detected in aggregates during anaerobic copper treatment (Zuily et al., 2022). LeuB (3-isopropylmalate dehydrogenase, EC 1.1.1.85) has been detected in protein aggregates when the gene was overexpressed in *E. coli* (Śmigiel et al., 2022) and in yeast deficient in the shock protein Hsp70 of type-Ssa (Amm, Kawan, and Wolf, 2016).

Decreased intracellular amino acid concentrations may result in numerous deleterious effects on bacterial physiology. These could include deficiencies in protein and other cell structure synthesis (e.g., cell wall), metabolism of various cell pathway intermediates, cellular buffering, and osmotolerance, among several others (Xiao et al., 2017; Nelson and Cox, 2017; Aliashkevich, Alvarez, and Cava, 2018; Idress et al., 2018). Interestingly, this metabolic condition could be somewhat beneficial; under physiological conditions of anaerobic amino acid limitation, the efflux system pump CusCFBA is induced to protect iron-sulfur cluster proteins on *E. coli* cells exposed to Cu(I) (Fung et al., 2013).

We also observed the accumulation of certain amino acids after metal(loid) exposure. For instance, serine accumulates after Au(III), Cd(II), Hg(II), and Te(IV) treatments, with the latter inducing the highest accumulation (4.3-fold) (**Figure 4C**). Similarly, Cd(II) treatment also resulted in an accumulation of glutamic acid and valine. The excess of intracellular serine is toxic for *E. coli* as it inhibits the biosynthesis of isoleucine and aromatic amino acids (Hama et al., 1990; Tazuya-Murayama et al., 2006), disrupts cell division (Zangh and Newman, 2008), and generates the misincorporation of serine instead of alanine in peptidoglycan crosslinks (Zangh, El-Hajj, and Newman, 2010). Serine excess is removed to the extracellular medium by SdaC or can be deaminated to pyruvate and ammonia by SdaA, SdaB, TdcG, TdcB, or Thr/Ser dehydrogenase IlvA (Shizuta et al. 1969; Su, Lang, and Newman, 1989; Su and Newman, 1991; Burman et al. 2004; Borchert and Downs, 2018; Kriner and Subramaniam, 2019). Mutants lacking these proteins did not display significant changes in fitness (CG z-scores) after soft-metal(loid) exposure and only IlvA was detected in the aggregated material produced by Te(IV) (**Table S1**). IlvA was also aggregated in treatments that did not induce serine accumulation, so it may not solely explain the phenomena observed with Te(IV). Our data do not suggest that serine accumulation is a direct result of protein aggregation. Altogether, our results demonstrate that soft-metal(loid)s result in the aggregation of proteins required for amino acid biosynthesis, which in turn leads to perturbations in the intracellular levels of amino acids.

In summary, we demonstrate that proteins can be aggregated in a ROS-independent manner by soft-metal(loid)s, mainly affecting proteins involved in anabolic pathways. Protein aggregation might be primarily induced at the translational level by Ag(I), Au(III), Hg(II), and Te(IV), or at the post-translational level by Cd(II) and As(III). Owing to their hydrophobic and disordered properties, initial protein aggregates might result in unspecific sequestration of other proteins, leading to secondary loss of function (Mogk et al., 2018). Regardless of the mechanism, it can be expected that amino acid biosynthesis is hindered as a cellular response under our experimental conditions, also impacting the tRNA charging process, ribosome stalling and limited translation, induction of the stringent response, and altered synthesis of various metabolites that use amino acids as precursors (Traxler et al., 2008; Nedialkova and Leidel, 2015; Wang et al., 2020). The fact that we did not observe a direct correlation between protein aggregation and cell viability suggests that soft-metal(loid) toxicity is a multifactorial phenomenon, an observation further confirmed by the wide variety of gene deletions that can lead to decreased fitness upon soft-metal(loid) challenges. These various targets could explain why these types of metal(loid)s are effective biocides.

## 5. Author Contributions

Conceptualization, F.A.C., F.A.A., C.M.M, J.M.S and C.C.V.; Methodology, F.A.C., F.A.A. and C.C.V.; Formal analysis, F.A.C., J.S.P and B.P.; Investigation, F.A.C., M.P.S, R.A.L, C.M.M, C.A.C., B.P. and E.H.M; Resources, C.C.V.; Writing Draft F.A.C., F.A.A., J.M.S, C.M.M, C.C.V, J.S.P, B.P. and E.H.M.; Visualization, F.A.C. and C.A.C.; Project Administration, F.A.A. and C.C.V.; Supervision, F.A.A. and C.C.V.; Funding Acquisition, F.A.A. and C.C.V.

## 7 Acknowledgments

We thank Dr. Roberto Molina-Quiroz (CECS, Valdivia, Chile) and Dr. James Imlay (Department of Microbiology, University of Illinois Urbana-Champaign) for critically reading the manuscript.

